# A spatial analysis of childhood cancer and industrial air pollution in a metropolitan area of Colombia

**DOI:** 10.1101/639773

**Authors:** Ana Maria Valbuena-Garcia, Laura Andrea Rodriguez-Villamizar, Claudia Janeth Uribe Pérez, Feisar Enrique Moreno Corzo, Rafael Gustavo Ortiz Martinez

## Abstract

**Background:** Air pollutants are considered carcinogenic to humans. In some European countries, an association with childhood cancer in children has been established. This phenomenon has not been addressed in Latin America, despite the spatial variability of air pollutants that may limit the extrapolation of the results to other geographical areas.

**Objective:** To conduct a spatial analysis of the relationship between childhood cancer and air pollution from industrial sources in a metropolitan area of Colombia.

**Methods:** Incident cases of childhood cancers were obtained from the Population-based Cancer Registry of the Bucaramanga Metropolitan Area (2000-2015). Local and focused cluster tests were used for the detection of spatial clusters and the Poisson multivariable model was used to evaluate the combined effects of spatial variables.

**Results:** The Kulldorff’s focused test found a significant spatial cluster (p=0.001) around one industrial agglomerate and the multivariable model results suggests that the distance effect is modified by the directional effect of the wind.

**Conclusion:** A spatial cluster of incident cases of childhood cancer occurred in the municipality of Bucaramanga. Our finding supports the hypothesis that childhood cancer might be related with industrial air pollution exposure in a Latin American city.

## Introduction

Childhood cancer (CC), defined here as cancer in children aged 0-14 years, is considered a group of rare diseases. Nevertheless, about one new case of CC is diagnosed in the world every two minutes [1] and it is the leading cause of non-accidental death for this age group [2]. Worldwide, approximately 215,000 cancers are diagnosed annually in children younger than 15 years [2], and in Latin America and the Caribbean 17,500 children are newly diagnosed every year. It is estimated that 84% of all cases of CC in the world occur in low-income and middle-income countries, where 90% of the child population live [3].

Knowledge regarding the etiology of CC is limited. Some authors argue that environmental factors can be the cause of 85-96% of all cancers in childhood [4] [5] [6] [7]. In recent years, studies conducted in England, Spain and the United States found an association between CC, especially leukemia and CNS tumors, and air pollution exposure [8] [9] [10]. Air pollution is considered the world’s largest single environmental health risk as the World Health Organization (WHO, 2016) affirmed that 92% of the world population lives in places where air quality levels are poor [11].

During 2013, the International Agency for Research on Cancer (IARC) found that there is sufficient evidence about the carcinogenicity of outdoor air pollution in general and fine particulate matter (PM_2,5_) [12]. Research published in Europe and North America, have found association between CC and proximity to industrial sources that emit pollutants into the air [5][9][10][13][14]; also, since the beginning of the 20th century, geographic clusters of cancer have been identified, mainly for leukemia [15]. The mixture of pollutants varies widely in space and time, reflecting the heterogeneity of their sources, the influence of climate, atmospheric transformations, and other factors, and therefore health effects might also differ widely in space and time and cannot be generalized.

In Colombia, a developing country located at the north extreme of South America, there are no exact estimates of the incidence of CC since 90% of the regions of the country do not have cancer registries, which limits making truthful national statistics [16]. Population-based registries with validated high quality data by the IARC are located only in the metropolitan areas of Cali, Manizales, Pasto and Bucaramanga [17]. In Colombia, outdoor air pollution is considered one of the three most hazardous environmental exposures [18] and annual mean concentrations of particulate matter in capital cities exceeds the ones reported for most European and US cities [19]. Previous studies in Colombia have identified adverse effects of ambient air pollutants on respiratory and cardiovascular mortality and morbidity [20,21].

Industrial sources are important contributors of outdoor air pollution and location of industrial facilities is often close to residential areas [22]. To our knowledge there are no studies in Colombia or other Latin American country that evaluate the geographic location of incident cases of CC with proximity to industrial sources of air pollution, despite the contribution of industrial emissions to outdoor air pollution in developing countries. Therefore, the aim of this study was to assess the spatial relationship between CC and air pollution from industrial sources in the metropolitan area of Bucaramanga, Colombia, during 2000-2015.

## Methods

### Study area and population

Colombia is the fourth largest country in South America located at the top corner of the continent. The study area was the Bucaramanga Metropolitan Area (BMA) that it is located in the Northeast Andean region of Colombia with an extension of approximately 1,479 km^2^ covering four municipalities: Bucaramanga, Floridablanca, Giron, and Piedecuesta. The study population were the children under 15 years old living in BMA; according to the National Administrative Department of Statistics (DANE for its acronym in Spanish) for the year 2015, the total population in BMA was 1,122,961 people and the population under 15 years was 258,097 children.

### Data Sources

The study used secondary retrospective data available in different regional and national institutions: The Population-based Cancer Registry of the Bucaramanga Metropolitan Area (RPC-AMB for its acronym in Spanish) was the source for cancer cases data, The DANE was the source for estimations of population and geographical data at small geographical level (census sector), the Corporation for the Defense of the Bucaramanga Plateau (CDMB for its acronym in Spanish) was the source for the identification and location of industrial facilities emitting air pollutants to the air, and planning office of each municipality was the source for the socioeconomic information at small area level.

#### Childhood cancer data

The CC data were obtained from the RPC-AMB, which is one of the four Population-based Cancer Registry of regions of Colombia considered by the IARC as high quality. Incident confirmed cases of CC diagnosed from January 1, 2000 to December 31, 2015 were included in the study. The information obtained for CC cases were: the type of cancer, sex, date of birth, date of diagnosis, and place of residence at the time of diagnosis. In the RPC-AMB and for the present research, it is considered a resident who is verified as having been residing 6 months or more in the BMA before the first time they were diagnosed with CC.

#### Population data and geographical data

The population data for the children’s population in the BMA were taken from the official DANE population estimates based on the 2005 national census. The projected infant population under 15 years was used as the population at risk to calculate CC rates during the study period. The DANE population estimates information is available with free access with a minimum level of geographic disaggregation up to block level that is added in an ascending way at the level of sections, sectors, communes, urban/rural areas and municipalities. For the analysis, the census sectors were used as geographical grouping unit (average population of each census sector is 1857 children under 15 years old); these geographical units have a unique identification number assigned by DANE and that was kept as the identification number for each unit of analysis.

The geographic coordinates (latitude and longitude) of each centroid of the geographic units (census sector) were estimated using the estimation method with weighting by population location and therefore the centroids are not geographical centers of polygons necessarily. For the analysis of proximity to sources of industrial pollution, the distance and direction between each “putative source” and the centroids of the census sectors were calculated using distance (in meters) and angle (in geodetic degrees) calculation tool of ArcGIS^®^ [23]. The map of the BMA with resolution at the block and sector census level was obtained from the Geoportal cartographic data tool of DANE [24] and the spatial data were created in ArcGIS^®^ using the projection of Colombia in mode Custom Azimuth Equidistant and Datum WGS 1984.

#### Data sources of industrial sources and socioeconomic status

The industrial facilities emitting air pollutants to the air were identified with the fixed sources inventory existing in the local environmental authority, the CDMB. There were 32 industrial facilities that were located spatially and displayed four identifiable industrial areas around and within BMA. The central point (centroid) of each identified industrial area was used as the location of the “putative source of industrial pollution” and was calculated as the geographic center of the area, georeferenced in latitude and longitude coordinates.

The socioeconomic strata (SES) is a DANE classification of the socioeconomic resources of a residential place [25] and was used as proxy of socioeconomic status. The predominant SES of each census sector was assigned according to the SES of the neighborhoods that make up each census sector. This information was obtained from the planning office of each municipality.

### Data analysis

#### Exploratory analysis

Crude rates (CR) of annual and cumulative CC incidence were calculated by census sector. The DANE population estimates for each geographical unit were used as population denominators of the annual and accumulated rates. Standardized rates were calculated by age and sex and their 95% confidence intervals, using the direct method and as a standard population, the population of Colombia was used by five-year groups according to the 2005 DANE Census. To reduce the heterogeneity in estimating the risk of CC, the calculation of a standardized morbidity ratio (SMR) of the incident CC was made using a Bayesian smoothing technique. Crude rates, standardized rates and Bayesian SMR were calculated in Stata 15^®^ and then visualized on choropleth maps using ArcGIS 12^®^.

#### Hypothesis tests

A cancer spatial cluster is defined as a region with a statistically significant excess in the number of cancer cases that occurs [26]. We used the Kulldorff circular space-time scanning test to identify spatial and spatio-temporal local clusters of CC. Three different hypothesis focused tests were used to identify potential spatial clusters of CC around the four identified industrial pollution areas. Based on the spatio-temporal parameters, retrospective space-time scanning analysis was applied to identify the geographic areas and time periods of potential clusters. The Kulldorff ‘s circular spatial scanning test in its focused version having as reference grid or “putative source” the coordinates of the centroid of each industrial area identified in the BMA; this test was implemented using the SaTSCan 9.1^®^ software, with a Poisson probability distribution model, scanning for high rates and with a maximum cluster size of 25% the population at risk [27]. The Stone’s test was used to evaluate wheather the risk of the disease is altered with the distance to the putative source; this test was implemented using the “DCluster” programming package encoded in the R software [28]. Finally, the Lawson’s directional score test was used to evaluate the variation in direction of the clusters using a score statistic for a single parameter that has a Chi2 distribution with one degree of freedom [29]; the score calculation was implemented in a Microsoft Excel^®^ spreadsheet.

#### Multivariable models

For the estimation of multivariable regression models with spatial data, we used the approach proposed by Lawson [30]. In the basic or null Poisson model, the log of the expected cases based on the number of populations by geographic area is used as an offset. Subsequently, the spatial functions (distance, direction and their interactions) are included in the multivariable model; to model direction, the functions of sine (longitude) and cosine (latitude) were used. These multivariable generalized linear models (GLM) for CC were constructed separately for each putative source of industrial pollution. Finally, the effects of the spatial functions were adjusted by the SES of the census sectors. The Akaike information criterion (AIC) and the deviance of the potential models were used as the parameters for model selection. The residuals of the models were estimated, and the spatial autocorrelation of the models was evaluated using the global Moran’s index. These analyzes will be performed using Stata 15^®^.

#### Missclassification bias and sensitivity analyses

A sub-sample of the CC were contacted by the RPC-AMB in order to assess the quality of the residential address at the time of diagnosis. We inquired about the address at the time of diagnosis and mobility in the two years prior to the time of diagnosis. With this information, the proportion of misclassification in the subsample was estimated to estimate the potential misclassification of the exposure in the study.

Sensitivity analysis was carried out for the local and focused cluster Kulldorf’s test. In both cases, the methodology was to execute the analysis with six circular search windows with different parameters, with the following specifications: radius of 0.5 km, 1 km, and 3 km and 5%, 10% and 25% of population, Poisson distribution, non-overlap parameter and level of significance of ≤0.05. Conducting multiple analyses using different search windows may strengthen findings and provide confidence about the detected CC clusters.

### Ethical considerations

Ethical approval for this study was obtained from the ethics committee of the Industrial University of Santander. The information received from the RPC-AMB did not contain identifying information about participants. The data was analyzed aggregated and not at the individual level, the georeferencing corresponds to the count of cases of CC for each census sector and it is not the exact georeferencing of the place of residence of the cases. For the analysis of misclassification where a subsample was recontacted, the recontact was managed with the caregiver of the minor through the attending physician. Informed consent was obtained from all participants of this subsample.

## Results

### Descriptive analysis of childhood cancer

From 2000 to 2015, 679 children under 15 years of age with a median age of 6 years (Interquartile range [IQR] 3 – 11 years) were diagnosed with some subtype of CC in BMA and 43.89% were girls. The median annual CC was 44 cases (IQR 38,5 – 46 cases). The two municipalities with the highest proportion of cases were Bucaramanga (54.05%) and Floridablanca (25.77%). The most commonly diagnosed types of cancer were: 1) leukemia, myeloproliferative diseases and myelodysplastic diseases (38.29%); 2) neoplasms of the central nervous system (CNS) and intracranial and intraspinal neoplasms (16.05%); and 3) lymphomas and reticuloendothelial neoplasms (13.4%). The overall CR of incident CC for BMA over the study period was 158.12 cases per million in children aged under 15 years; the overall annual CR increased from a minimum of 103.53 in 2000 to a maximum of 195.62 in 2013. The overall direct sex and age standardized rate (SASR) for BMA was 158.92 cases per million (IC 95%: 147.18 – 171.34); according to the 95% confidence interval (95% CI), the overall SASR of the municipality of Bucaramanga was significantly higher than that of Giron and Piedecuesta. Table 1 shows the overall crude and age-standardized incidence rates by sex for AMB and each municipality.

**Table 1.** Overall incidence rates for BMA and its municipalities by sex, 2000-2015

| Region | Crude rate |  | Age-standardized rate |  |  |  |
| --- | --- | --- | --- | --- | --- | --- |
|  | per million children |  | per million children under 15 years old |  |  |  |
|  | under 15 years old |  |  |  |  |  |
|  | *Female* | *Male* | *Female* | *95% CI* | *Male* | *95% CI* |
| **AMB** | 141.72 | 173.85 | 142.35 | (126.60 –<br>159.43) | 174.76 | (157.64 –<br>193.23) |
| **Bucaramanga** | 156.44 | 203.30 | 157.01 | (133.24 –<br>183.79) | 203.97 | (177.31 –<br>233.51) |
| **Floridablanca** | 140.89 | 190.69 | 142.32 | (111.33 –<br>179.24) | 193.29 | (157.41 –<br>234.87) |
| **Giron** | 110.23 | 103.31 | 110.33 | (77.25 –<br>152.76) | 103.29 | (71.94 –<br>143.66) |
| **Piedecuesta** | 120.27 | 110.30 | 120.67 | (83.06 –<br>169.46) | 109.80 | (75.09 –<br>155.04) |

BMA is divided into 151 census sectors, in 121 (80.13%) there was at least one case of CC during 2000-2015. The overall CR in the census sector varied from a maximum of 866.35 to a minimum of 36.72 cases per million, with a median of 177.04 (IQR: 103.1 – 259.7). The SASR were between 33.78 (95% CI: 4.09 – 125.89) and 795.67 cases per million (95% CI: 20.14 – 4650.31), with a median of 179.18 (IQR: 103.2 – 263.6). A census sector located in the northwest of the municipality of Bucaramanga, had the overall SASR significantly higher than the overall SASR of BMA. The Bayesian SMR in the census sectors ranged from 0.63 to 1.49 (median: 1.02, IQR: 0.87 – 1.14). Figure 1 show the choropleth maps of the SASR of CC for BMA over the study period.

**Figure 1.**
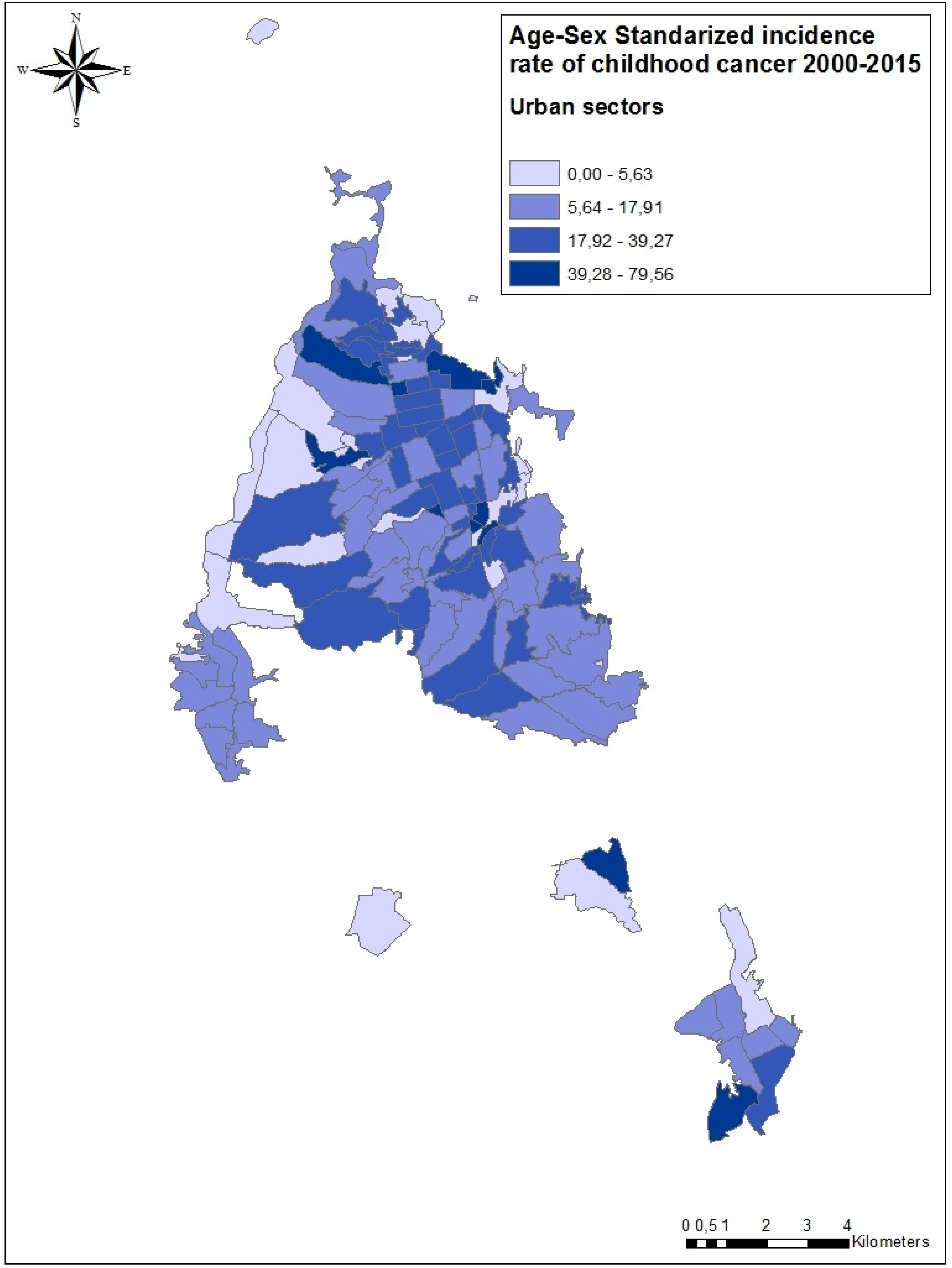
Age and sex standardized rates of childhood cancer in the urban area of the municipalities of the BMA during 2000 to 2015.

### Descriptive analysis of industrial sources

The BMA inventory included a total of 38 fixed industrial sources, 78.95% were constituted before the year 2000, 7.89% in 2004 and 13.16% between 2007 and 2009. Four main areas of industrial facilities agglomerations were identified, to which the centroid and the coordinates were calculated; these four industrial areas and their centroids were used as a source of exposure for subsequent analyzes. The first agglomerate was referenced as industrial area A with latitude 7.138530546 and longitude −73.13197221 and located in the census sector 680011000000000306. The second referenced as industrial area B with latitude of 7.095961661 and longitude −73.16810226 and located in the census sector 683071000000000001. The third area is the denominated C with latitude 7.081772454 and longitude −73.15087541 and located in the census sector 680011000000001196. The fourth industrial area, the D has latitude 7.06152597 and longitude −73.09061454, located in the census sector 682761000000000009 (Fig 2).

**Figure 2.**
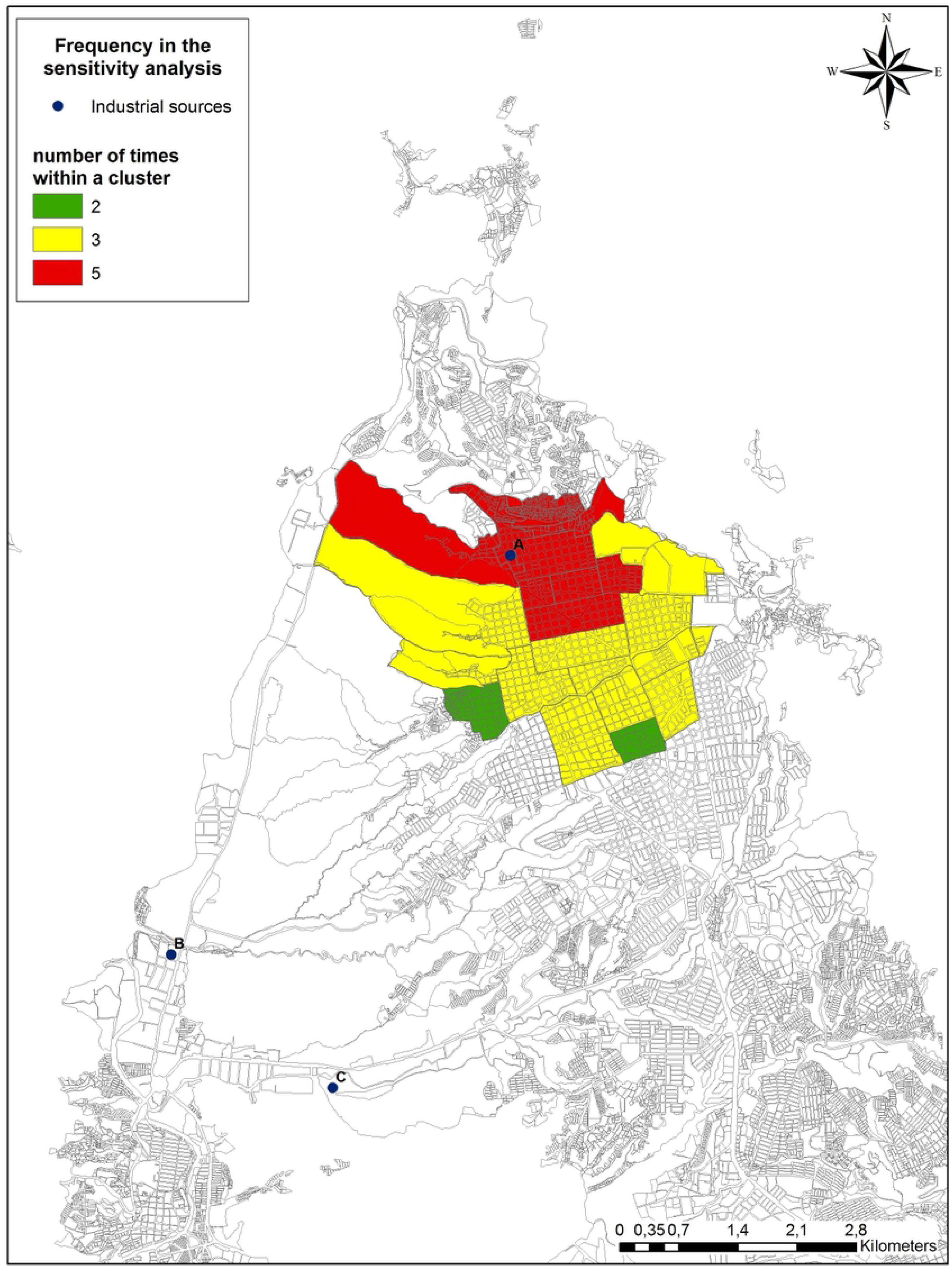
Census sectors detected within the results of the six analyzes of focused childhood cancer clusters through different search windows, Bucaramanga, Colombia, 2000-2015

### Hypothesis tests

Of the total of cases, 598 (88.07%) had data for georeferencing. The Kulldorff circular scan local test identified a spatio-temporal cluster of statistically significant of CC cases (p<0.001), with a radius of 1.64 km and estimated relative risk of 1.79 in the municipality of Bucaramanga; furthermore, it identified a space-time cluster of CC cases covering 18 census sectors in the municipality of Bucaramanga, the high-risk period was the year 2009 and estimated relative risk of 6.21 (p:0.02). Around the industrial agglomerate A, the focused cluster test of Kulldorff found a significant cluster (p:0.001) with a radius of 2.65 km and estimated relative risk of 1.41 in the municipality of Bucaramanga, being coincident with the cluster identified by the local test. No significant clusters were found around the other three points of industrial area with this test. Stone’s test did not detect significant clusters for the points of industrial agglomerates. Lawson’s directional score was significant from industrial areas A, B and C. Table 2 summarizes the results of the three focused tests located for the four industrial areas.

**Table 2.** Results of focused hypothesis tests for clusters of childhood cancer in proximity to industrial areas in BMA, 2000-2015

| Industrial areas | Kulldorf's circular spatial |  | Stone's test for |  | Lawson's directional |  |
| --- | --- | --- | --- | --- | --- | --- |
|  | scan test |  | distance decline |  | score test |  |
|  | Statistic | p-value | Statistic | p-value | Statistic | p-value |
| **A** | 12.27 | <0.001 | 2.44 | 0.09 | 6.82 | 0.01 |
| **B** | No clusters found |  | 1.02 | 0.89 | 13.17 | <0.001 |
| **C** | 0.002 | 0.86 | 1.45 | 0.25 | 6.93 | 0.01 |
| **D** | 2.11 | 0.08 | 1.19 | 0.29 | 0.94 | 0.33 |

### Multivariable Analysis

The Moran coefficient was 0.059 (p <0.001) for the total number of CC events, which means that there was a low spatial autocorrelation among the census sectors. The distribution of the SMR as outcome variable of the multivariable model (mean: 1.09, variance: 1.00) showed to be appropriated for the proposed Poisson model.

The results of the multivariable Poisson models support a consistent significant inverse effect of the distance to the industrial area A on the SMR (Table 3). The model that evaluated only the directional effect had a significant result for the positive effect of the longitude, suggesting a dominant effect toward the East from the industrial area A. The model that evaluates the combined effect of distance and direction suggested a significant effect by decreasing the distance and directional towards the East from A. When the interaction terms between the distance and the address parameters were added to the model and adjusted by the SES, the model improved the deviance suggesting the best model quality. In this model the inverse effect of the distance to industrial area A remained significant, and the statistical significance of the interaction terms suggest that the distance effect is modified by the directional effect (that is, the smaller the distance the stronger is the directional effect), the coefficients and signs of the interaction terms suggest that the predominant directionality is towards the SouthEast direction from A (Table 3).

**Table 3.**
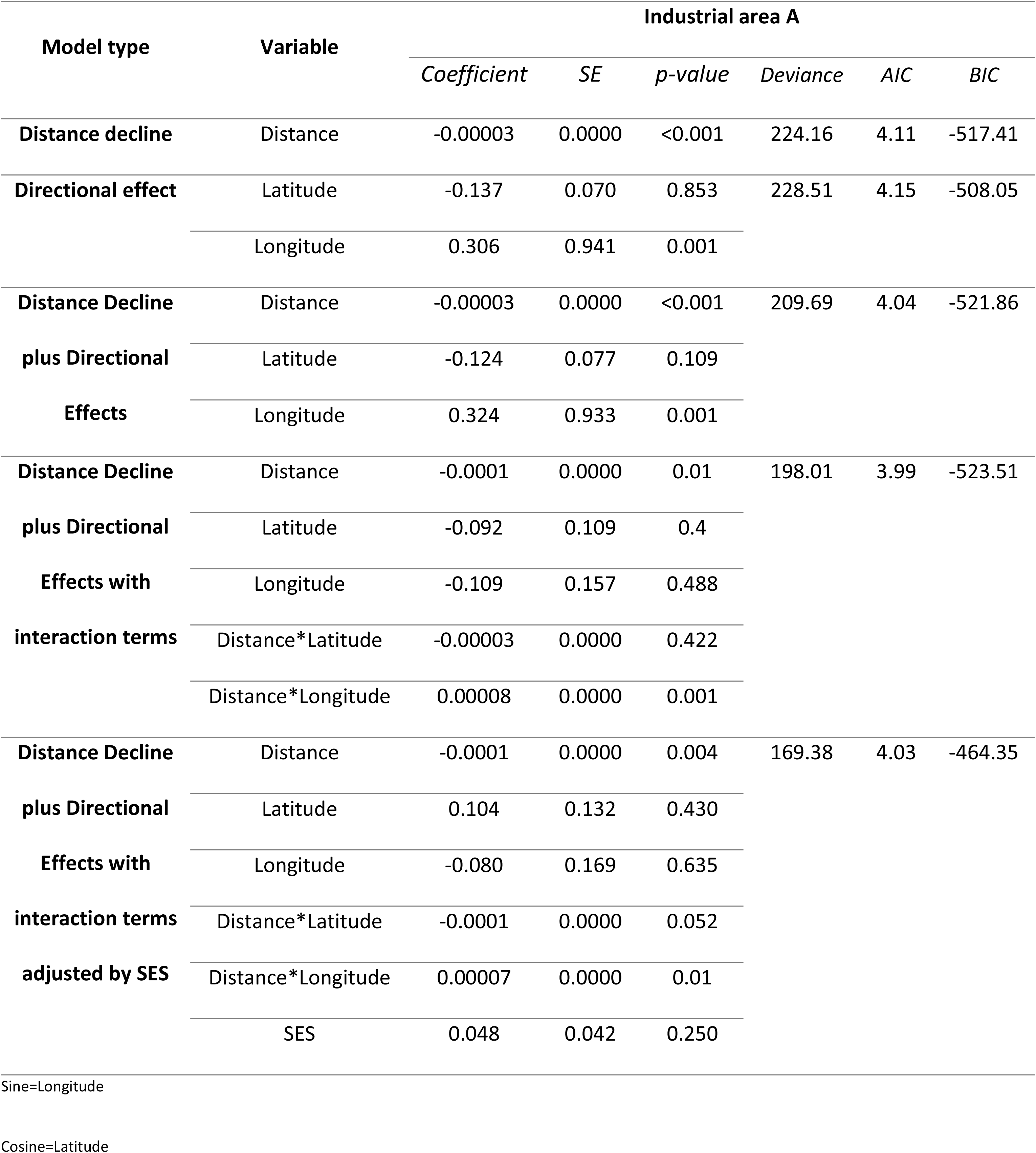
Poisson models assessing distance and directional effects of childhood cancer rates around the industrial area A, 2000-2015

In the model diagnosis, standardized Pearson, Deviance and Ascombe residuals fulfill the assumption of normal and mean distribution close to zero. Finally, the Moran coefficient for the residuals was 0.74 (p <0.001), which means that there is a low spatial autocorrelation of the residuals of the model.

For the industrial areas B, C and D the multivariable models adjusted by ESE that incorporated the terms of interaction between distance and directionality were also those that had the best criteria to be the best model and in no case the models had significant results for the effect of distance or directional effect.

### Sensitivity analysis

#### Misclassification analysis

A total of 40 CC cases were contacted. The median age was 6 years (IQR 2 – 9) and 86.84% were residents in neighborhoods of SES 1 to 3. Of the total, four (10%) had changed their homes in the year prior to diagnosis. Of the four cases with mobility in the last year prior to diagnosis, two had changed their housing within the same municipality, one between the municipalities of the BMA and one to a municipality outside the BMA. A total of 22 (55%) children resided in the same home since birth.

#### Sensitivity analysis for Kulldorff tests

In the local Kulldorf’s test, a cluster of CC cases with statistical significance was identified in the northwest zone of municipality of Bucaramanga, which was repeated in 5 of the 6 windows executed in the 1km circular window; 12 census sectors were identified that extend to 26 in the 3km window and that were equivalent to the analysis with a population threshold at risk of 5% and 25%, respectively. Figure 2 shows in red census sectors that were detected repeatedly within 5 clusters, in yellow those detected within 3 clusters and in green those detected within 2 clusters.

In the focused Kulldorf’s tests, taking as reference the industrial area A, in the six search windows a spatial cluster was identified, with two census sectors that were always present in the six analyzes executed, 680011000000000306 (Sector that corresponds to the area industrial A) and 680011000000000305. In the 0.5km circular window, two census sectors were identified that extend to 41 in the 3km window and that were equivalent to the analysis with a population threshold at risk of 5% and 25%, respectively.

## Discussion

The incidence rates of CC in the BMA between 2000 and 2015 are higher than the estimated world rates for the same period of time. The cumulative SASR 2000-2015 in the BMA was 1.5 times higher than the world rate reported by the World Cancer Report in 2014 that was 100 CC cases per million [31]. However, the cumulative SASR in the BMA was similar to the cumulative incidence rate of CC reported by the Population-based Cancer Registry of Cali during 2002-2011 [32]. The cancer registry of Cali is the oldest in Latin America and has international recognition for its high quality methods and data [33][34], therefore, the similarities in the incidence of CC and the frequency of presentation of the different types represent not only the quality and maturity of the RPC-AMB but also the consistency of having higher CC rates in different cities in Colombia.

The findings of this study suggest the presence of a CC cluster in proximity to the industrial area denominated A. These results were consistent across the descriptive, hypothesis testing and multivariable analysis findings, and remained statistically significant in the same area when different scan windows were used in the sensitivity analysis. The Lawson’s directional score and in the Poisson model that evaluated distance and directionality indicate that there is a spatial presentation of the cases that is mostly concentrated at southeast of industrial area A, finding that might be explained by the potential dispersion of air pollutants as the direction of the cluster is pointing towards the predominant direction of wind in the area [29].

The Stone’s test was the test that had no statistically significant findings for the industrial area A; this negative result might be explained by the fact that the Stone’s test is executed under the assumption that there is a decreasing trend in risk symmetrically around the radius as the distance to the source increases, and as shown in the multivariable analysis and sensitivity analysis (Fig 2) the risk is not symmetric but with direction towards southeast from A [35]. The statistically significant results in the industrial areas B and C for the directional score of the Lawson test seem to be isolated findings as they were not consistent with the multivariable analysis, where the interaction between distance and direction was evaluated as suggested by Lawson [36].

The spatial association found in this research should not be taken as a causal relationship between the industrial emissions of the industrial area A and the incident cases of the CC cluster in this area of the municipality of Bucaramanga. In this case, the relationship between exposure and outcome was modeled with respect to distance to the source and directionality of the wind, but the exposure to the pollutants emitted were measured. Likewise, the retrospective evaluation of the exhibition is difficult since it usually involves different changes during the study period [37].

When a cluster of a disease is detected the Centers for Disease Control and Prevention (CDC) proposes a four-stage approach, where the final stage includes proposing a formal etiological investigation [38] and it is the proposal that is suggested as the next step to this study. The increase of cases of IQ in this area can be due to multiple factors such as the proximity to exit routes of the city with high flow of heavy duty vehicles of diesel engines, which the exhaust of this type of engine is classified within the group 1 of carcinogens by IARC, which has been found associated with CC [39]. As also for the permanent transit of this type of automotive in this area, due to the loading and unloading of merchandise in the industrial sources that are in this industrial agglomerate. Likewise, in this area of industrial agglomerate the exhibition may not be related to air emissions but to the occupational exposure of parents who live near the workplace; there are several authors who have found a relationship between childhood cancer and maternal or paternal occupation [40] [41] [42] [43] [44].

### Consistency with previous studies

Different studies in Europe have found an association between exposure to air pollution and CC. Steffen et al. carried out a multicenter case control study in France and found an association between dwellings neighbouring a petrol station or a repair garage during childhood and the risk of childhood leukemia (OR 4.0, 95% CI 1.5 to 10.3) [45]. During 2009, a study with the same aim was also conducted in France and the findings supported the results and the associations established by Steffen [13].

In Spain several studies have been conducted assessing the association between spatial location of industrial pollution sources and CC. During 2015, Ramis et al. published a case-control study with spatial analysis, which included cases of leukemia, cancer of the CNS and non-Hodgkin’s lymphoma from five regions of Spain diagnosed from 1996 to 2011, and reported absence of any statistically significant cluster [46]. Ortega-García and cols., analyzed the spatial distribution of all incident cases of CC diagnosed in Spain during 1998 to 2015, using focused cluster tests and found a possible association between proximity to certain industries and CC [10]. Previous population case-control studies conducted in Spain by same research team reported an increased risk of Leukemia in children (OR: 1.31, 95% CI: 1.03-1.67) [5], and renal tumors (OR: 1.97, 95% CI: 1.13-3, 42) [47] associated with living at a distance less than 2.5 km from industrial facilities; increased risk of neuroblastomas associated with industrial facilities at 1 km (OR: 2.52; 95% CI: 1.20-5.30) and 2 km (OR: 1.99; 95% CI: 1.17-3.37) [48]; and increased risk of bone tumors in children under 15 years of age associated with living less than 3 km from industrial sources (OR: 2.33; 95% CI: 1.17-4.63) [49]. They found no excess risk for retinoblastoma, liver tumors, soft tissue sarcomas and germ cell tumors and living close to industrial areas [50].

### Strengths and limitations

The main strength of this study is the quality of the CC data and the period of 16 years included for the analysis. The cancer registries are considered the best source of information to measure cancer burden in a given time and region [15] [51], the RPC-AMB is considered high quality by IARC due to the integrity, punctuality and accuracy of the data provided, which translates into highly standardized and reliable data. Therefore, the classification of the disease is considered adequate for the cases included in the analyzes performed. In addition, it can be considered that the results of this research are not affected by the mobility of the population, since the misclassification analysis conducted did not found that the CC had a high proportion of migration and 55% of the cases in BAM lived in the same residence where they received the diagnosis since birth.

Another important strength to note is the use of multiple analytical approaches and tests used to evaluate the existence and characteristics of a potential CC cluster. The combination of descriptive, hypothesis testing, and multivariable analysis improved the quality of the analysis and gave consistency to the results.

The limitation of the pre-established shapes that the Kulldorff’s tests used in this study have been discussed in the literature [52][53][54][55]. These circular clusters in many cases generate that areas of the detected spatial cluster correspond to a false positive, since those areas with low risks are not a genuine part of the cluster [53][55]. To overcome this limitation this study includes other hypothesis test and multivariable analysis that include direction testing and their interaction with distance and results of the existence of a cluster identified by scanning tests remained consistent with additional information related to the direction of the cluster.

The correct classification of the exposure is a limitation and it is considered that to a certain extent all the studies evaluating the effects of exposure to air pollution inevitably have a degree of misclassification of exposure, since it is not feasible to conduct population–based studies in which each person uses a personal measuring device for long time [56]. In this study, the magnitude of the exposure was probably determined not only by the distance to the industrial areas identified in the BMA, but also for quantity or air pollutant emissions and the predominant wind direction in the industrial areas. Although the predominant wind direction was taken into account for directional tests, the variability in time of the wind direction was not included. Other important limitation of this study is that exposure was based on proximity to industrial facilities only and did not include any characterization or measurement of the pollutants emitted to the air by the industrial sources. Therefore, proximity to industrial facilities was used in this study as proxy of industrial air pollution exposure.

## Conclusions

The results of this study suggest the presence of a spatial cluster of incident cases of CC during 2000-2015 in proximity to an industrial area in the municipality of Bucaramanga. Given the ecological design of this research this finding cannot be assumed as a causal relationship, but results of this study generate new research hypotheses to be evaluated in further research in order to contribute to have a better understanding of the environmental exposures related to the high incidence of CC in Bucaramanga and Colombia.

## Authors’ contributions

AMVG participated in the design of the research, performed the non-spatial and spatial statistical analyses and drafted the manuscript. LARV participated in the design of the research, review of the analyzes, coordination of the study. FEMC and RGOM contributed with support the geographical data processing, visualization and analysis. CJUP participated with obtaining and reviewing data on childhood cancer and subsamples. LARV, FEMC, RGOM, CJUP have critically revised the article and all authors have approved the final manuscript.

## REFERENCES

1. WHO. Globocan 2012 [Internet]. Available from: http://globocan.iarc.fr

2. IARC. International Childhood Cancer Day: Much remains to be done to fight childhood cancer. 2016;(February):1–2. Available from: https://www.iarc.fr/wp-content/uploads/2018/07/pr241_E.pdf

3. Magrath I, Steliarova-Foucher E, Epelman S, Ribeiro RC, Harif M, Li CK, et al. Paediatric cancer in low-income and middle-income countries. Vol. 14, The Lancet Oncology. 2013. p. e104–16.

4. Ferrís Tortajada J, Ortega García JA, Marco Macián A, García Castell J. Medio ambiente y cáncer pediátrico. An Pediatría [Internet]. 2004;61(1):42–50. Available from: https://linkinghub.elsevier.com/retrieve/pii/S1695403304783526

5. García-Pérez J, López-Abente G, Gómez-Barroso D, Morales-Piga A, Pardo Romaguera E, Tamayo I, et al. Childhood leukemia and residential proximity to industrial and urban sites. Environ Res [Internet]. 2015 Jul;140:542–53. Available from: https://linkinghub.elsevier.com/retrieve/pii/S0013935115001607

6. Gouveia-vigeant T, Tickner J. Toxic chemicals and childhood cancer: A review of the evidence. A Publication of the Lowell Centre for Sustainable Production. University of Massachusetts Lowel. 2003.

7. Narod S, Stiller C, Lenoir G. An estimate of the heritable fraction of childhood cancer. Br J Cancer [Internet]. 1991 Jun;63(6):993–9. Available from: http://www.nature.com/doifinder/10.1038/bjc.1991.216

8. Knox E. Childhood cancers and atmospheric carcinogens. J Epidemiol Community Heal [Internet]. 2005 Feb 1;59(2):101–5. Available from: http://jech.bmj.com/cgi/doi/10.1136/jech.2004.021675

9. Filippini T, Heck JE, Malagoli C, Giovane C Del, Vinceti M. A Review and Meta-Analysis of Outdoor Air Pollution and Risk of Childhood Leukemia. J Environ Sci Heal Part C. 2015;33(1):36–66.

10. Ortega-García JA, López-Hernández FA, Cárceles-Álvarez A, Fuster-Soler JL, Sotomayor DI, Ramis R. Childhood cancer in small geographical areas and proximity to air-polluting industries. Environ Res. 2017;156:63–73.

11. World Health Organization. Ambient Air Pollution: A global assessment of exposure and burden of disease. 2016.

12. IARC/WHO. Outdoor Air Pollution. In: Monographs on the Evaluation of Carcinogenic Risks to Human. Lyon; 2016.

13. Brosselin P, Rudant J, Orsi L, Leverger G, Baruchel A, Bertrand Y, et al. Acute childhood leukaemia and residence next to petrol stations and automotive repair garages: the ESCALE study (SFCE). Occup Environ Med [Internet]. 2009 Sep 1;66(9):598–606. Available from: http://oem.bmj.com/cgi/doi/10.1136/oem.2008.042432

14. Lavigne É, Bélair M-A, Do MT, Stieb DM, Hystad P, van Donkelaar A, et al. Maternal exposure to ambient air pollution and risk of early childhood cancers: A population-based study in Ontario, Canada. Environ Int. 2017 Mar;100:139–47.

15. Goodman M, Lakind JS, Fagliano JA, Lash TL, Wiemels JL, Winn DM, et al. Cancer cluster investigations: Review of the past and proposals for the future. Vol. 11, International Journal of Environmental Research and Public Health. 2014. p. 1479–99.

16. Bravo LE. Estimating the incidence and mortality of cancer in Colombia: What are the best data for public policies? Colomb Med. 2016;47(2):71–3.

17. Forman D, Bray F, Brewster D., Gombe-Mbalawa C, Kohler B, Piñeros M, et al. Cancer incidence in five continents, vol. X. In WHO Press; 2014. Available from: http://ci5.iarc.fr/CI5I-X/old/vol10/CI5vol10.pdf

18. Planeación Nacional. Consejo Nacional de Política Económica y Social (CONPES) 3550. Republica de colombia. Departamento de planeación. 2008.

19. Instituto de Hidrología Meteorología y Estudios Ambientales – IDEAM. Informe del Estado de la Calidad del Aire en Colombia 2011-2015. 179 p.

20. Blanco-Becerra LC, Miranda-Soberanis V, Hernández-Cadena L, Barraza-Villarreal A, Junger W, Hurtado-Díaz M, et al. Effect of particulate matter less than 10μm (PM10) on mortality in Bogota, Colombia: A time-series analysis, 1998-2006. Salud Publica Mex. 2014.

21. Rodríguez-Villamizar LA, Rojas-Roa NY, Fernández-Niño JA. Short-term joint effects of ambient air pollutants on emergency department visits for respiratory and circulatory diseases in Colombia, 2011–2014. Environ Pollut. 2019 May;248:380–7.

22. Larsen B. Cost of Environmental Damage: A Socio-Economic an Environmental Health Risk Assessment [Internet]. 2004. Available from: http://www.bvsde.paho.org/texcom/cd050996/larsen.pdf

23. ESRI. ArcGIS Desktop: Release 10.2. Redlands CA. 2013.

24. DANE Departamento Administrativo Nacional de Estadisticas. Geoportal [Internet]. Available from: https://geoportal.dane.gov.co

25. Departamento Administrativo Nacional de Estadística (DANE) de Colombia. Estratificación Socioeconómica. Estadísticas por Tema. 2016.

26. National Center for Environmental Health. CDC – Cancer Clusters – Home. [cited 2017 Feb 21]; Available from: https://www.cdc.gov/nceh/clusters/

27. Kulldorff M, Information Management Services. SaTScanTM v8.0: software for the spatial and space-time scan statistics [Internet]. 2009. Available from: http://www.satscan.org/

28. R Core Team. R Software [Internet]. R: A Language and Environment for Statistical Computing. Vienna, Austria; 2013. Available from: http://www.r-project.org/

29. Lawson AB. On the Analysis of Mortality Events Associated with a Prespecified Fixed Point. J R Stat Soc Ser A. 1993;156(3):363–77.

30. Lawson A. Statistical Methods in Spatial Epidemiology. Wiley Series in Probability and Statistics. 2006. 424 p.

31. Stewart BW, Wild CP. World cancer report 2014. World Health Organization. Lyon, France; 2014.

32. Bravo LE, García LS, Collazos P, Aristizabal P, Ramirez O. Descriptive epidemiology of childhood cancer in Cali, Colombia 1977-2011. Colomb Med. 2013;44(3):155–64.

33. Correa P. The Cali Cancer Registry an example for Latin America. Colomb medica (Cali, Colomb. 2012 Oct;43(4):244–5.

34. García LS, Bravo LE, Collazos P, Ramírez O, Carrascal E, Nuñez M, et al. Cali cancer registry methods. Colomb Med [Internet]. 2018;49:109–20. Available from: http://colombiamedica.univalle.edu.co/index.php/comedica/article/view/3853

35. Stone RA. Investigations of excess environmental risks around putative sources: Statistical problems and a proposed test. Stat Med. 1988;7(6):649–60.

36. Lawson AB. The statistical analysis of points events associated with a fixed point. University of St Andrews; 1991.

37. Lawson A, Banerjee S, Haining R, Ugarte L. Handbook of Spatial Epidemiology. 2016. 704 p.

38. CDC. Guidelines for investigating clusters of health events. Vol. 30, Mortality and Morbidity Weekly Report. 1990.

39. Boothe VL, Boehmer TK, Wendel AM, Yip FY. Residential traffic exposure and childhood leukemia: A systematic review and meta-analysis. Am J Prev Med. 2014;46(4):413–22.

40. Feychting M, Plato N, Nise G, Ahlbom A. Paternal occupational exposures and childhood cancer. Environ Health Perspect. 2001;109(2):193–6.

41. Castro-Jiménez MÁ, Orozco-Vargas LC. Parental exposure to carcinogens and risk for childhood acute lymphoblastic leukemia, Colombia, 2000-2005. Prev Chronic Dis. 2011;8(5):A106.

42. Park AS, Ritz B, Ling C, Cockburn M, Heck JE. Exposure to ambient dichloromethane in pregnancy and infancy from industrial sources and childhood cancers in California. Int J Hyg Environ Health. 2017 Oct;220(7):1133–40.

43. Spycher BD, Lupatsch JE, Huss A, Rischewski J, Schindera C, Spoerri A, et al. Parental occupational exposure to benzene and the risk of childhood cancer: A census-based cohort study. Environ Int. 2017 Nov;108:84–91.

44. Peters S, Glass DC, Greenop KR, Armstrong BK, Kirby M, Milne E, et al. Childhood brain tumours: associations with parental occupational exposure to solvents. Br J Cancer. 2014 Aug 24;111(5):998–1003.

45. Steffen C, Auclerc MF, Auvrignon A, Baruchel A, Kebaili K, Lambilliotte A, et al. Acute childhood leukaemia and environmental exposure to potential sources of benzene and other hydrocarbons; a case-control study. Occup Env Med. 2004;61(9):773–8.

46. Ramis R, Gómez-Barroso D, Tamayo I, García-Pérez J, Morales A, Pardo Romaguera E, et al. Spatial analysis of childhood cancer: a case/control study. PLoS One. 2015;10(5):e0127273.

47. García-Pérez J, Morales-Piga A, Gómez J, Gómez-Barroso D, Tamayo-Uria I, Pardo Romaguera E, et al. Association between residential proximity to environmental pollution sources and childhood renal tumors. Environ Res. 2016;147:405–14.

48. García-Pérez J, Morales-Piga A, Gómez-Barroso D, Tamayo-Uria I, Pardo Romaguera E, Fernández-Navarro P, et al. Risk of neuroblastoma and residential proximity to industrial and urban sites: A case-control study. Environ Int. 2016;92:269–75.

49. García-Pérez J, Morales-Piga A, Gómez-Barroso D, Tamayo-Uria I, Pardo Romaguera E, López-Abente G, et al. Risk of bone tumors in children and residential proximity to industrial and urban areas: New findings from a case-control study. Sci Total Environ. 2017;579:1333–42.

50. García-Pérez J, Morales-Piga A, Gómez-Barroso D, Tamayo-Uria I, Pardo Romaguera E, López-Abente G, et al. Residential proximity to environmental pollution sources and risk of rare tumors in children. Environ Res. 2016;151:265–74.

51. Bray F, Znaor A, Cueva P, Korir A, Swaminathan R, Ullrich A, et al. Planning and Developing Population-Based Cancer Registration in Low-And Middle-Income Settings. Vol. 43, IARC Technical Publication. 2014.

52. Puett RC, Lawson AB, Clark AB, Hebert JR, Kulldorff M. Power evaluation of focused cluster tests. Environ Ecol Stat. 2010;17(3):303–16.

53. Grubesic TH, Wei R, Murray AT. Spatial Clustering Overview and Comparison: Accuracy, Sensitivity, and Computational Expense. Ann Assoc Am Geogr. 2014;

54. Duczmal L, Kulldorff M, Huang L. Evaluation of Spatial Scan Statistics for Irregularly Shaped Clusters. J Comput Graph Stat. 2006 Jun;15(2):428–42.

55. Jacquez GM. Cluster morphology analysis. Spat Spatiotemporal Epidemiol. 2009;1(1):19–29.

56. Oudin A, Forsberg B, Strömgren M, Beelen R, Modig L. Impact of Residential Mobility on Exposure Assessment in Longitudinal Air Pollution Studies: A Sensitivity Analysis within the ESCAPE Project. Sci World J. 2012;2012:1–5.

